# Out-of-distribution prediction with disentangled representations for single-cell RNA sequencing data

**DOI:** 10.1101/2021.09.01.458535

**Authors:** Mohammad Lotfollahi, Leander Dony, Harshita Agarwala, Fabian Theis

**Author notes:** Equal contribution (Mohammad Lotfollahi, Leander Dony, Harshita Agarwala). Correspondence to: Mohammad Lotfollahi < >.

## Abstract

Learning robust representations can help uncover underlying biological variation in scRNA-seq data. Disentangled representation learning is one approach to obtain such informative as well as interpretable representations. Here, we learn disentangled representations of scRNA-seq data using *β* variational autoencoder (*β*-VAE) and apply the model for out-of-distribution (OOD) prediction. We demonstrate accurate gene expression predictions for cell-types absent from training in a perturbation and a developmental dataset. We further show that *β*-VAE outperforms a state-of-the-art disentanglement method for scRNA-seq in OOD prediction while achieving better disentanglement performance.

## 1. Introduction

In disentanglement learning, a single latent dimension is linked to a single generative feature, while being relatively invariant to changes in other features (Ridgeway, 2016; Bengio et al., 2013; Chen et al., 2016). Inversely, by manipulating values of a single dimension in the latent space, only a single generative feature is perturbed. Such representations allow for more interpretable latent spaces. This is particularly relevant for scRNA-seq data, where finding a biologically interpretable representation is desired. The characteristics of a disentangled representation also allow for Out of Distribution (OOD) prediction. In OOD prediction, the gene expression of a cell-type absent from training in the desired condition is extrapolated. This could apply to an unmeasured cell-type after a perturbation or as part of a developmental trajectory. Recent models have addressed OOD predictions for scRNA-seq in context of perturbation and disease. (Lotfollahi et al., 2019b;a). Although these models aim to provide a meaningful representations of the data, they do not learn a disentangled representation. The work proposed in (Lopez et al., 2018) is a step towards interpreting scRNA-seq with disentangled representations. The authors propose a VAE regularized with d-variable Hilbert-Schmidt Independence Criterion (dHSIC), which improves hypothesis testing by removing information corresponding to nuisance variables related to quality control measures. However, in addition to VAE parameters, the dHSIC framework requires additional hyper-parameters to optimize the HSIC loss, making the optimization problem even harder. The authors also did not provide insight into the identity of individual learned dimensions for single-cell, thus adding no additional interpretability to current methods. To address these problems we propose to use a *β*-VAE model (Higgins et al., 2017; Burgess et al., 2018), a fully unsupervised model for disentanglement learning. We apply *β*-VAE on scRNA-seq data and show that the model successfully decomposes data into major interpretable generating factors such as cell-types and perturbations. We further demonstrate that obtained interpretable factors can be exploited for OOD predictions. Finally, we illustrate that *β*-VAE outperforms dHSIC in both disentanglement learning and OOD prediction.

## 2. Methods

A modified Variational Autoencoder (VAE) (Kingma & Welling, 2013; Kingma et al., 2014; Rezende et al., 2014) was employed for disentanglement learning as proposed previously by (Higgins et al., 2017; Burgess et al., 2018). It includes a linearly increasing controlling capacity *C* such that the KL divergence term is driven towards *C*. This allows more information to flow through the latent space, thus encouraging disentanglement. The deviation of the KL divergence loss from *C* is penalized by *β*. This model is referred to as ‘*β*-VAE’ model. Here the two available tuning parameters were *β* and *C*.

The second approach restricts the search space for the approximated posterior in a VAE by kernel-based measures of independence (Lopez et al., 2018). In particular it uses the d-variable Hilbert-Schmidt Independence Criterion (dHSIC) (Gretton et al., 2005; Pfister et al., 2016) to enforce independence between the latent representations and arbitrary nuisance factors. This was modified to enforce independence between the different dimensions of latent space. By scaling and adding this to the original VAE objective function, a new regularisation criteria was created. Therefore, by penalizing the dHSIC kernel value for sampled latent space, the model makes the dimensions independent of each other which in turn encourages disentanglement. This model is referred to as ‘dHSIC-VAE’ model. Here the two parameters were *β*, weight for KL divergence loss and *γ*, weight for dHSIC kernel value.

To identify suitable hyper-parameters for the models, we systematically ran grid-searches and selected a set of hyperparameters for each model that provided both good disentanglement as well as good reconstruction performance on the validation data.

To quantify disentanglement, we used the Disentanglement Score proposed by (Higgins et al., 2017). It relies on the main assumption that a few of the generative factors are conditionally independent and also interpretable. To apply the metric, data points which shared a single feature (e.g. same cell-type) were randomly picked. They were mapped to the latent space and their differences to each other were calculated. If the independence and interpretability properties hold for the learnt representations, there should be less variance in the inferred latent dimension that correspond to the fixed feature (cell-type in this case). A linear classifier was then used to identify this fixed feature by classifying the difference value in latent spaces and report the accuracy as the final disentanglement metric score. Smaller variance in the latent unit corresponding to the target factor will make the classifier more accurate, resulting in a higher score.

## 3. Results

### 3.1. Disentanglement learning enable predicting cellular responses

To compare disentanglement performance, we trained both *β*-VAE and dHSIC on a peripheral blood mononuclear cell (PBMC) scRNA-seq data. This dataset contains, 13944 cells, 14053 genes and 12 different cell-types in both control and *IFN-β* stimulated conditions (Kang et al., 2018). Both models learnt to disentangle two features ‘condition’ and ‘cell-type’. The dHSIC model also gave promising results, however not better than the *β*-VAE. Figure 1 compares the latent representations and values of KL divergence loss. It can be seen from the KL loss plots in the figure, that only the units that were active earlier in the training phase are disentangled. Additionally, the units that gain high value of KL loss later in the training, are not disentangling with ‘condition’ or ‘cell-type’ features. They could be associated with some other unknown generative factors. Table 1 compares the disentanglement scores. It can be seen that the scores are very close for both models. However, the *β*-VAE model performs slightly better than the dHSIC model.

**Figure 1.**
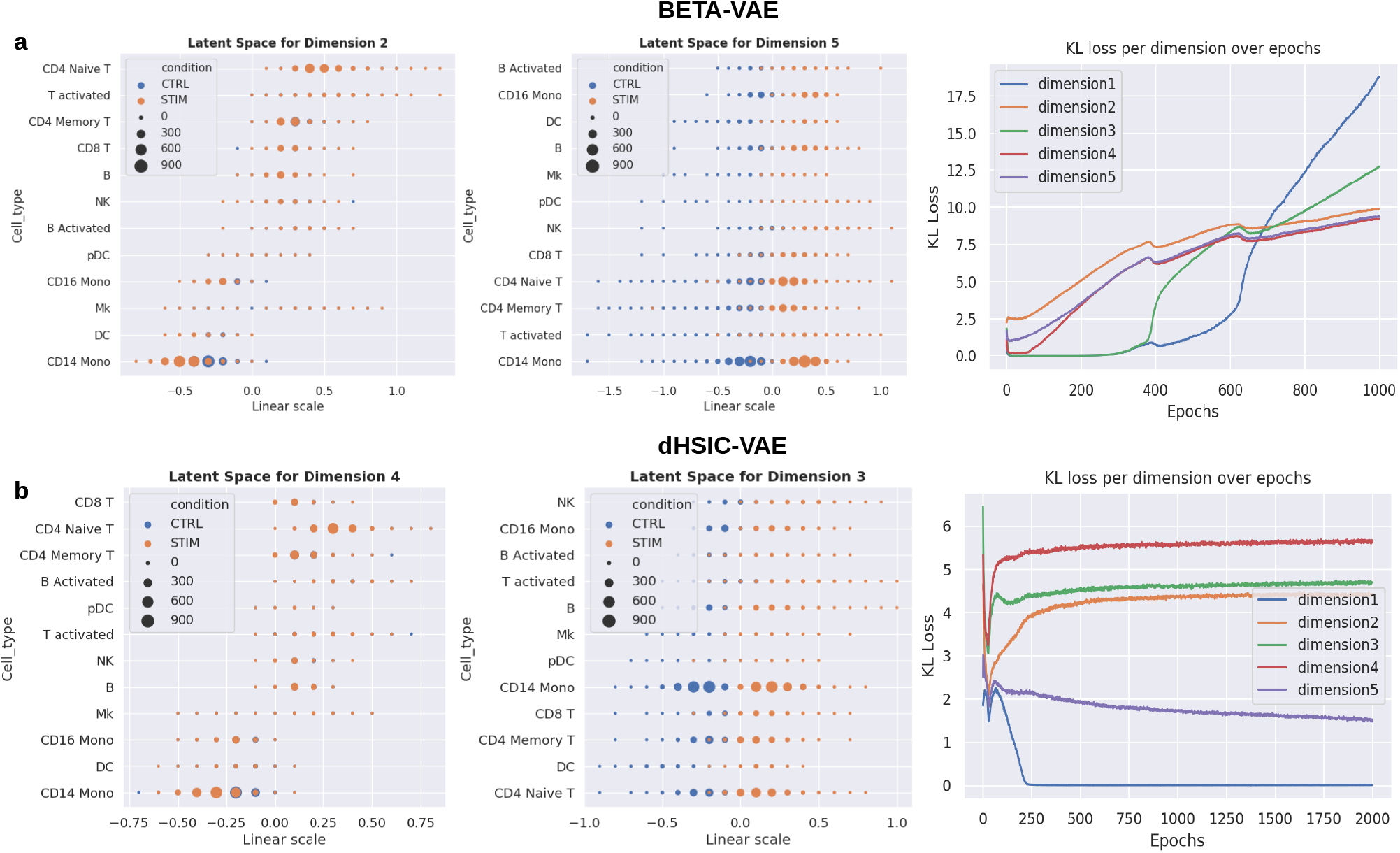
Comparison between latent representations of *β*-VAE (*β*=5 and *C*=30) and dHSIC (*β*=1 and *γ*=50) models in the Kang dataset. **(a)** shows units that disentangle ‘cell-type’ and ‘condition’ best in *β*-VAE along with KL divergence plot. Similarly, **(b)** shows the units for dHSIC model.

**Table 1.**
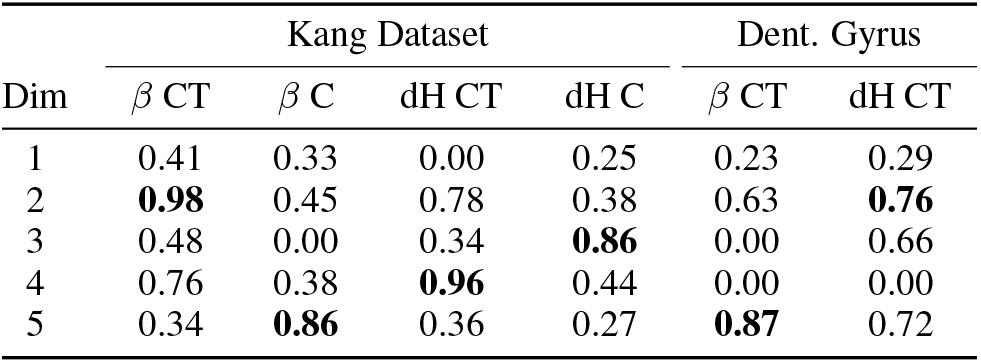
Disentanglement Scores (best dimension in bold). *β*-VAE model on Kang data: *β*=5, *C*=30; dHSIC model on Kang data: *β*=1, *γ*=50. *β*-VAE model on Dent. Gyrus data: *β*=50, *C*=30; dHSIC model on Dent. Gyrus data: *β*=1, *γ*=50. *β*: *β*-VAE model; dH: dHSIC model; CT: cell-type disentanglement; C: condition disentanglement; Dim: latent dimension.

Next, we sought to predict OOD ‘*CD4 Naive T*’ cells. We excluded both control and stimulated ‘*CD4 Naive T*’ cells during the training phase. After training, we identified the latent dimensions encoding ‘cell-type’ feature. By linearly interpolating this dimension, we could start from one celltype and then generate new cells that would vary in only ‘cell-type’ feature, recovering the held-out cells. We took a stimulated *B*-cell (source cell) and mapped it to the latent space. Then, we manipulated the values in the dimension that disentangled ‘cell-type’. This way we could recover stimulated ‘*CD4 Naive T*’ cell (target cell). As this dimension encoded ‘cell-type’ feature only, we kept the perturbation feature invariant and changed only ‘cell-type’. Similarly, we recovered control *‘CD4 Naive T’* from a control ‘*B*’ cell-type using the same dimension that encoded ‘cell-type’ feature. Figure (2) compares OOD prediction from both the *β*-VAE model and dHSIC. *β*-VAE outperformed dHSIC by achieving more accurate predictions (Figure 2, left column). The PCA plot shows that the predicted cells in the *β*-VAE are closer to the real cells when compared to dHSIC.

**Figure 2.**
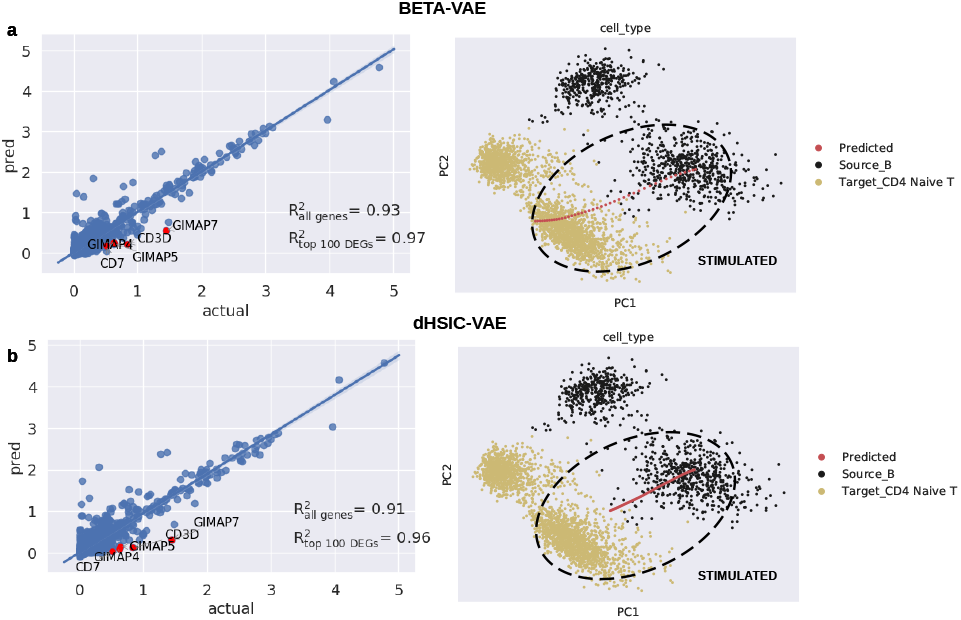
**(a-b)** OOD prediction comparison between *β*-VAE (*β*=20 and *C*=30) model and dHSIC (*β*=1 and *γ*=50) models. *R*^2^ denote the squared Pearson correlation of mean gene expression between predicted and real cells. Red dots denote top 5 DEGs in target cells. The PCA plot shows the target cell (‘*CD4 Naive T*’), source cells (‘*B*’) and the newly predicted cells.

### 3.2. Disentanglement learning allows recovery of missing cell-type from developmental trajectory

We further evaluated disentanglement performance of both models on a dataset from mouse dentate gyrus (Hochgerner et al., 2018). The data consists of 25,919 genes across 2,930 cells forming multiple lineages. The cells are grouped into 14 clusters by graph-based clustering. In the Dentate Gyrus dataset, both models learnt to disentangle cell-types. The dHSIC model also gave promising results, however not better than the *β*-VAE. Figure 3 compares the latent representations and value of KL loss for both the models. The units that were active initially and had the highest KL divergence showed the most disentanglement. Table 1 compares the disentanglement scores. It can be seen that *β*-VAE had higher scores than dHSIC model.

**Figure 3.**
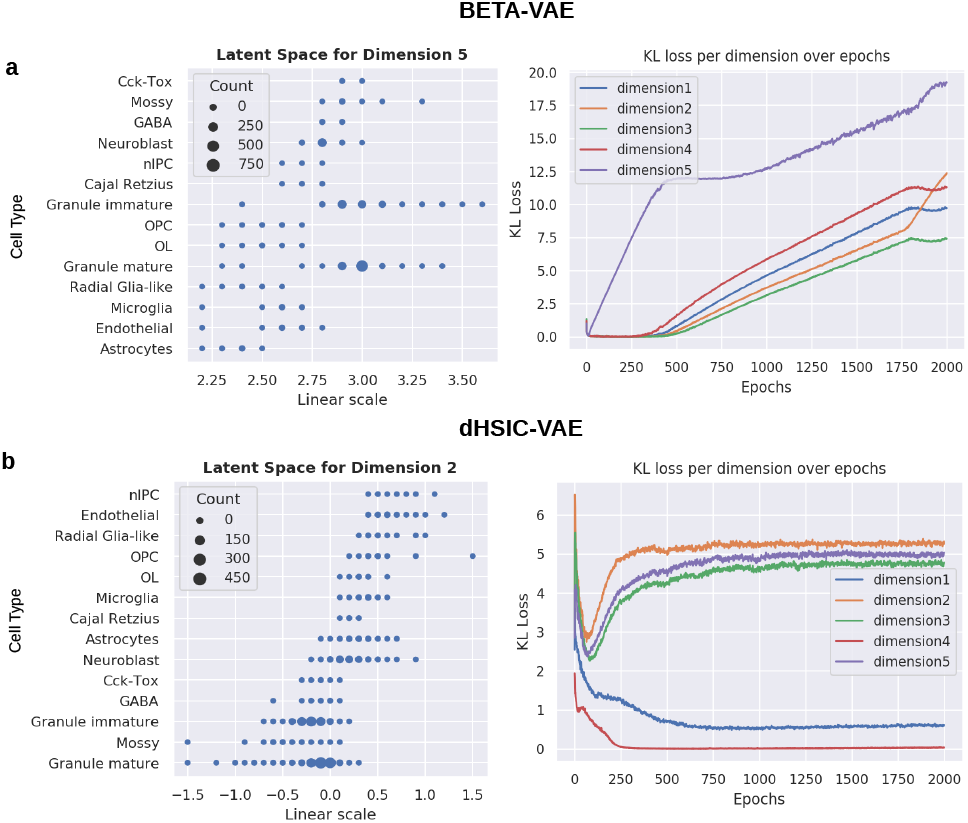
Comparison between latent representations of *β*-VAE (*β*=50 and *C*=30) and dHSIC (*β*=1 and *γ*=50) models in the Dentate Gyrus dataset. **(a)** shows units that disentangle ‘cell type’ best in *β*-VAE along with KL divergence plot. Similarly, **(b)** illustrates the unit for dHSIC model.

We hypothesised the cell type dimension could also capture the order of development and thus represent a developmental trajectory. To test this, we performed velocity analysis using scVelo (Bergen et al., 2019). A dominating developmental trajectory can be seen starting from ‘*nIPC*’ to ‘*Granule immature*’ via ‘*Neuroblast*’ (Figure 4). It shows that ‘*Neuroblast*’ develops into ‘*Granule immature*’ cells. We excluded ‘*Granule immature*’ cells from the training data and attempted to recover them using ‘*Neuroblast*’ cells. We sought to find latent dimensions that would encode the development trajectories of these cells. We tested recovery of ‘*Granule immature*’ through linear interpolation starting from ‘*Neuroblast*’ for all dimensions. The dimension that recovered ‘*Granule immature*’ was the one that disentangled cell types.

**Figure 4.**
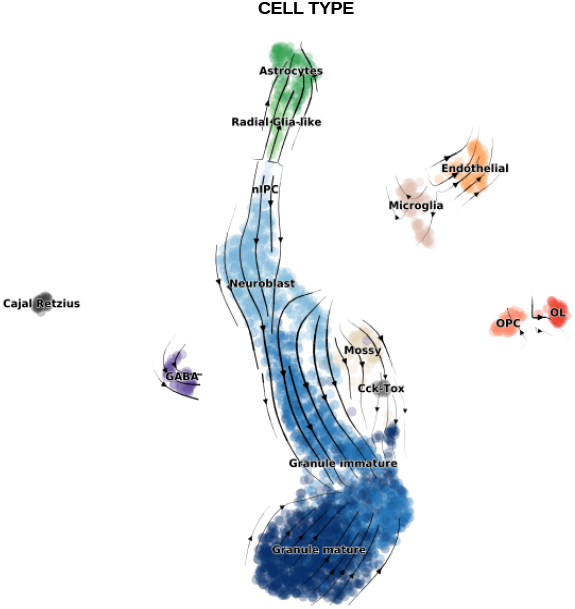
RNA Velocity on Dentate Gyrus dataset.

We observe *β*-VAE achieved more accurate results (Figure 5 left column) while having smoother transition between source and target cell types (Figure 5 right column). This observation confirms that *β*-VAE accurately captured continuous transition between cell types in a developmental trajectory.

**Figure 5.**
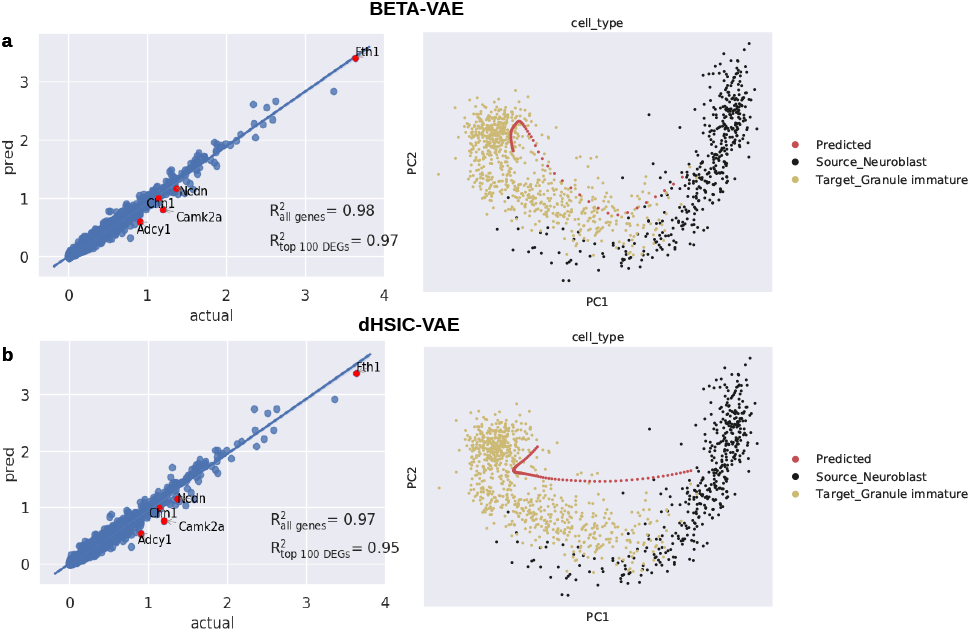
**(a-b)**, OOD prediction comparison in Dentate Gyrus Dataset between *β*-VAE (*β*=100 and *C*=30) and dHSIC (*β*=1 and *γ*=50) models. *R*^2^ denote the squared Pearson correlation of mean gene expression between predicted and real cells. Red dots denote top 5 DEGs in target cells. The PCA plot shows the target cell (‘*Neuroblast*’), source cells (‘*Granule immature*’) and the newly predicted cells.

## 4. Conclusion

We demonstrated that disentanglement representation learning provides interpretable factors for downstream tasks. We exemplified this by leveraging these factors to predict gene expression of cell types not seen in training data after a perturbation and also during brain development. We further illustrated that the *β*-VAE model achieves better feature disentanglement and prediction than the state-of-the-art dHSIC method on single-cell data. We envision that disentanglement learning on single-cell data can provide more interpretable representations leading to better understating of the underlying biology and cellular heterogeneity. The code for the model and the accompanying data can be obtained from https://bit.ly/3bRLpzL

